# CD9 co-operation with syndecan-1 is required for a major staphylococcal adhesion pathway

**DOI:** 10.1101/2023.01.17.524294

**Authors:** Luke R. Green, Rahaf Issa, Fawzyah Albaldi, Lucy Urwin, Henna Khalid, Claire E. Turner, Barbara Ciani, Lynda J. Partridge, Peter N. Monk

## Abstract

**Objectives:** Epithelial colonisation is a critical first step in bacterial pathogenesis. *Staphylococcus aureus* can utilise several host factors to associate with cells, including α5β1 integrin and heparan sulphate proteoglycans, such as the syndecans. Here, we demonstrate that a partner protein of both integrins and syndecans, the host membrane adapter protein tetraspanin CD9, is essential for syndecan-mediated staphylococcal adhesion. Fibronectin is also essential in this process while integrins are only critical for post-adhesion entry into human epithelial cells.

**Methods and Results:** Treatment of epithelial cells with CD9-derived peptide or heparin caused significant reductions in staphylococcal adherence, dependent on both CD9 and syndecan-1. Exogenous fibronectin caused a CD9-dependent increase in staphylococcal adhesion whereas blockade of β1 integrins did not affect adhesion but did reduce the subsequent internalisation of adhered bacteria. CD9 disruption or deletion increased β1 integrin-mediated internalisation, suggesting that CD9 coordinates sequential staphylococcal adhesion and internalisation.

**Conclusions:** CD9 controls staphylococcal adhesion through syndecan-1, using a mechanism that likely requires CD9-mediated syndecan organisation to correctly display fibronectin at the host cell surface. We propose that CD9-derived peptides or heparin analogues could be developed as anti-adhesion treatments to inhibit the initial stages of staphylococcal pathogenesis.

## Introduction

*Staphylococcus aureus* is an opportunistic pathogen and a common causative agent of community and nosocomial infections. Infections result in a myriad of clinical outcomes ranging from superficial skin infections to systemic infections including pneumonia, endocarditis and osteomyelitis (Tong et al., 2015). Furthermore, the rapid acquisition of various antimicrobial resistance mechanisms by *S. aureus* worldwide (Stryjewski and Corey, 2014) makes the study of this organism, and development of potential therapeutics, critical for healthcare. *S. aureus* has adapted a range of adhesins for adherence to host cell surface molecules, allowing for efficient downstream entry into cells (Josse et al., 2017). Several of these adhesins bind to the extracellular matrix (ECM) protein, fibronectin (Fn). The canonical cellular staphylococcal adhesion and internalization pathway utilises Fn-binding proteins (FnBPs) to bind to Fn using a zipper-like mechanism (Sinha et al., 1999). On the host cell surface, Fn binds to α5β1 integrins through an RGD motif, leading to the proposal that Fn acts as a bridge between the host and the pathogen. The FnBP-Fn-α5β1 complex is thought to be central to *S. aureus* adhesion and internalization (Josse et al., 2017).

*S. aureus* can also utilise heparan sulphate proteoglycans (HSPGs), such as the syndecans (SDC), to adhere and gain entry into host cells (Hess et al., 2006). Many bacteria utilise the glycosaminoglycan (GAG) heparan sulphate (HS) moiety of proteoglycans (PG), using these negatively charged polysaccharides as binding sites for clusters of positively charged amino acids within FnBPs (Chen et al., 2008). SDC-1 in particular has been associated with *S. aureus* adherence to intestinal, lung and corneal epithelial cell lines, since interactions were blocked using HS mimics such as heparin (Hess et al., 2006; García et al., 2016; Rajas et al., 2017). SDC-1^−/−^ mice were also more resistant to corneal infection by *S. aureus,* however, no specific binding of *S. aureus* to SDC-1 was demonstrated, suggesting that it is not directly involved in attachment (Hayashida et al., 2011). SDC can also bind Fn through HepII domains (Gong et al., 2008) and have been shown to modulate Fn fibrillogenesis in an integrin-dependent manner (Yang and Friedl, 2016). This has led to suggestions that SDC, integrins and Fn collaborate in the formation of binding sites on host cells that are exploited by many pathogenic bacteria, for example *Streptococcus pneumoniae* (Jinno et al., 2020). However, the molecular details of these interactions are currently unknown.

The tetraspanins are a superfamily of transmembrane proteins whose major function is to organise various proteins at the cell surface into dynamic structured ‘islands’, known as tetraspanin-enriched microdomains (TEMs) (Hemler, 2005). There are 33 human tetraspanins with conserved structural motifs; four transmembrane domains, a small (EC1) and large (EC2) extracellular loop and generally short N-and C-terminal domains. The crystal structures of CD9 and CD81 demonstrate a reverse cone-like structure creating a lipid binding pocket within the transmembrane domains. Lipid interactions with this pocket are thought to change the conformation of the tetraspanin between an ‘open’ and ‘closed’ state (Zimmerman et al., 2016; Umeda et al., 2020). In the open state, the EC2 domain is available for partner protein interactions that mediate the involvement of TEMs in a wide variety of cellular functions (Yang et al., 2020). For example, the tetraspanins have been shown to organise partner proteins into ‘adhesive platforms’ to allow leukocyte adhesion to endothelial cells (Barreiro et al., 2008). Partner proteins can include integrins, HSPGs and immunoglobulin superfamily members (Umeda et al., 2020), many of which are important in host cell adherence and entry of microbial pathogens. The tetraspanins have been implicated in a number of different viral and bacterial infections (Florin and Lang, 2018; Karam et al., 2020). We have previously demonstrated that tetraspanin blockade, using antibodies or recombinant EC2 domains, can significantly reduce Gram-negative and Gram-positive bacterial adherence to epithelial cells (Green et al., 2011). We further showed that short CD9 EC2-derived peptides were able to significantly reduce *Staphylococcus aureus* adherence to keratinocytes and to an engineered model of human skin (Ventress et al., 2016).

In the present study, we demonstrate for the first time that the tetraspanin, CD9, is a critical component of an SDC-1/Fn adhesion network exploited by *S. aureus* as a primary means of adhesion to human epithelial cells. Staphylococcal adherence was significantly reduced by both a CD9 EC2-derived peptide and by soluble GAG analogues (heparins), but no additive effect was observed in combination. We show that proteoglycan SDC-1 is essential for these inhibitory effects. Addition of exogenous Fn to epithelial cells increases staphylococcal adherence but this is completely abrogated by CD9 EC2-derived peptide. Integrin α5β1 blockade has no effect on bacterial adherence but CD9 appears to inhibit α5β1-mediated staphylococcal internalization, suggesting that CD9 coordinates adhesion and internalization. Taken together, our study demonstrates the critical importance of CD9 during Fn-mediated staphylococcal adherence and highlights the potential of the tetraspanin-derived peptides and heparins as anti-adhesive therapeutics for bacterial infections.

## Materials and Methods

### Strains and bacterial growth conditions

*S. aureus* strains, SH1000 and SH1000 Δ*fnb,* were kindly provided by Simon Foster (University of Sheffield, UK). MRSA isolates were retrieved from skin infections of patients at Northern General Hospital, Sheffield and kindly provided by Sue Whittaker. Liquid cultures were grown in Luria broth (LB; Oxoid, Ltd., UK) microaerobically at 37°C in a humidified atmosphere with constant agitation. Solid cultures were grown on LB agar (Oxoid) overnight at 37°C. Freshly grown plates were used to inoculate all liquid cultures.

### Cell culture

Wild-type (WT) and CD9 knockout (^−/−^) A549 human lung epithelial cells were a gift from David Blake (Fort Lewis College, Colorado, USA) (Blake et al., 2018) while Tspan15^−/−^ A549 cells were obtained from Mike Tomlinson (University of Birmingham, UK). HaCaT cells, a human keratinocyte cell line, were supplied by Cell Lines Service (CLS GmbH, Eppelheim, Germany). IPEC-J2 cells, a porcine intestinal epithelial cell line, were supplied by DSMZ-German Collection of Microorganisms and Cell Culture (Braunschweig, Germany). The above cell lines were maintained in Dulbecco’s Modified Eagle’s Media (DMEM; ThermoFisher Scientific, Massachusetts, USA) and 10% heat-inactivated foetal calf serum (FCS; Bioscience, UK). HCE-2 (ATCC, Virginia, USA), a human corneal epithelial cell line, was cultured in keratinocyte serum-free media (KSFM; ThermoFisher Scientific) supplemented with hydrocortisone, insulin, epidermal growth factor and bovine pituitary extract.

Primary normal human tonsillar keratinocytes (NTK) were a gift from Craig Murdoch and Helen Colley (School of Clinical Dentistry, University of Sheffield, UK). NTK were isolated from palatine tonsils collected from patients during routine tonsillectomies at the Royal Hallamshire Hospital, Sheffield Teaching Hospitals NHS Foundation Trust with written, informed consent (UK National Research Ethical Committee approval number 09/H1308/66) and cultured in flavin and adenine-enriched medium as previously described (Grayson et al., 2018). For experiments, cells were differentially trypsinised to remove the irradiated 3T3 fibroblast feeder layer, and NTK seeded on surfaces coated with 100 μg/ml type IV human collagen (Merck) in defined keratinocyte serum-free medium (ThermoFisher Scientific) supplemented with Y-27632 (Abcam, Cambridgeshire, UK).

### Peptides, antibodies and GAGs

Peptides were synthesised using solid phase Fmoc chemistry (Genscript, New Jersey, USA). The CD9 EC2-derived peptide, 800C (DEPQRETLKAIHYALN), was designed using the 15 residue segment of the second α-helix from the EC2 domain, previously demonstrated to inhibit staphylococcal interactions with human cells (Ventress et al., 2016). A homologous EC2-derived peptide was designed within the related tetraspanin CD81 (CD81 P1: DANNAKAVVKTFHETLD). Scrambled peptides were randomly generated from the CD9 (QEALKYNRAEETPLDIH) and CD81 sequences (ADTDALVNKFTKHANEV). Fluorescently-labelled peptides were synthesised with a 6-carboxyfluorescein moiety at the N-terminus. Integrin RGD peptides (GRGDS & SDGRG), heparinase I/III, chondroitinase ABC and fondaparinux were obtained from Merck, Germany. Unfractionated heparin (Leo) and dalteparin were obtained from the Royal Hallamshire Hospital Pharmacy, UK. Heparan sulphate (HS), chondroitin sulphate (CS) and dermatan sulphate (DS) were supplied by Iduron, UK. Mouse anti-human SDC-1 IgG1 (B-A38; Santa Cruz Biotechnology, Texas, USA), mouse anti-human SDC-4 IgG2a (5G9; Santa Cruz Biotechnology), rat anti-human β1 IgG1 (AIIB2; Merck), mouse anti-human β1 IgG1 (Lia1/2; GeneTex, California, USA), mouse anti-human α5 IgG_1_ (JBS5; Merck), mouse anti-human α5 IgG3 (P1D6; Merck), mouse IgG1 (JC1; in house), mouse IgG2a (02-6200; Thermofisher Scientific), mouse IgG3 (B10; Thermofisher Scientific) and rat IgG1 (14-4301-82; Thermofisher Scientific), mouse IgG1 ascites fluid (MOPC 21; Merck) were used as described. Pixatimod was kindly provided by Zucero Pty, Brisbane, Australia.

### SDC knockdown by shRNA

Short hairpin RNA (shRNA) targeting SDC-1, −4 or a nonsense control inserted into the lentiviral plasmid pLVTHM were obtained from Andreas Ludwig (RWTH Aachen University, Germany) (Pasqualon, 2015; Pasqualon et al., 2015). Sub-confluent HEK293T cells were co-transfected with 2.6 μg pMD2G (Addgene, USA), 7.4 μg psPAX2 (Addgene) and 10 μg pLVTHM with 10 μl jetPEI (Polyplus transfection, France) to produce recombinant lentiviruses. After 24 h, media was changed and the resulting lentivirus-containing supernatants were harvested after a further 48 h. For transient transfection, 2−10 WT cells were grown for 24 hours before replacement of 20% of the culture media with lentivirus-containing supernatant. Transduction efficiency was enhanced through addition of polybrene (Merck). GFP expression was used to assess the efficiency of transduction and flow cytometry was used to measure the expression of SDC1 and −4.

### Expression analysis

Tetraspanin and staphylococcal receptor expression was measured by flow cytometry. 1×10^6^ adherent cells were detached using cell dissociation buffer (ThermoFisher Scientific) and labelled with the relevant antibody at 4°C for 60 minutes. A fluorescein isothiocyanate (FITC) conjugated goat anti-mouse IgG antibody (F5387, Merck) was used for secondary labelling if required. Cells were fixed with 1% paraformaldehyde, analysed with an LSRII cytometer (Becton Dickinson, Oxford, UK) and analysed with FlowJo v10.0.7r2 software (BD).

### Fluorescent 800C binding

WT or CD9^−/−^ cells were seeded at 2.5×10 onto sterile 22mm glass coverslips and incubated overnight at 37°C with 8% CO_2_. Cells were washed with PBS, before addition of 200nM 6-carboxyfluorescein (FAM) tagged peptide or media alone for 30 minutes at 37°C. Cells were fixed with 2% PFA for 15 minutes before being washed twice. Coverslips were placed on slides with Vectashield Mounting Media with DAPI (Maravai LifeSciences, California, USA) and imaged using a Nikon Ti Eclipse microscope. Images were analysed using ImageJ and corrected total fluorescence intensity was calculated as follows: Total cell fluorescence = Integrated density - (area of selected cell or whole image x mean fluorescence of background).

### Infection assays

2×10^4^ cells were seeded into 96 well plates and cultured overnight. 5% bovine serum albumin (BSA; Merck) was added to cells for 60 minutes to reduce non-specific binding. Cells were washed with PBS and treated with peptide, antibodies or blocking reagents for a further 60 minutes. Cells were infected at a multiplicity of infection (MOI) of 50 for 60 minutes at 37°C with 5% CO_2_. Cells were washed four times with PBS and lysed with 2% saponin (Merck) for 30 minutes. Lysates were serially diluted, plated onto LB agar plates and allowed to grow overnight at 37°C. To further control for non-specific plastic binding, the number of bacteria bound to BSA blocked empty wells were subtracted from adherent and internalised bacteria. Bacterial adherence to treated cells was calculated as a percentage of bacterial adherence to untreated cells which was set at 100%.

For enumeration by microscopy, 1.5×10 WT cells were seeded onto glass coverslips in 24 well plates and cultured overnight. Cells were blocked, treated and infected as described above. Cells were washed four times with PBS after infection to remove unbound bacteria. Coverslips were fixed with a methanol:acetic acid (3:1) solution for 5 minutes. Fixed cells were washed with distilled water and stained with 10% Giemsa stain for 20 minutes. Coverslips were mounted and viewed by light microscopy. 100 cells were counted from various fields of view and scored for the number of infected cells and the total number of adhered bacteria.

### Gentamicin protection assays

Infection assays were carried out as described. After infection, cells were washed four times with PBS before immersion in cell media with 200 μg/ml gentamicin for one hour to eliminate extracellular bacteria. Wells containing no cells were used to ensure efficient killing of bacteria by the antibiotic. Cells were washed twice with PBS and lysed with 2% saponin for 30 minutes. Lysates were serially diluted and plated onto LB agar before overnight incubation at 37°C.

### Statistical analyses

All analyses were performed within GraphPad Prism version 9.0.0 (GraphPad Software Inc., USA). Significance was established at *p*<0.05, all data presented represents at least three independent experiments. Statistical considerations and specific analyses are described separately within each section. * specify significance to the untreated control unless otherwise specified; * *p*≤0.05, ** *p*≤0.01, *** *p*≤0.001.

## Results

### CD9-blockade reduces staphylococcal adherence to cells

Tetraspanins, HSPGs and β1 integrins have individually been demonstrated to be involved in staphylococcal adherence and internalization (Sinha et al., 1999; Hess et al., 2006; Green et al., 2011; Ventress et al., 2016). To investigate the relationship between these groups of molecules, staphylococcal adherence was measured by colony forming unit (CFU) after treatment of WT A549 or tetraspanin CD9^−/−^ A549 cells with a CD9-derived peptide (800C). CD9 was well expressed in WT cells but significantly reduced in CD9^−/−^ cells (Fig. S1A, B). 800C significantly reduced *S. aureus* SH1000 adherence to WT cells at 20nM (48.7±11.6%) but was slightly less effective at 2nM (29.3±9.2%; Fig. 1A). Pre-treatment with a scrambled 800C sequence peptide caused no significant reduction at concentrations up to 200nM (Fig. 1A). With untreated CD9^−/−^ cells, staphylococcal adherence was significantly reduced compared to untreated WT cells (41.5±11.4%; Fig. S1C) but treatment with 800C or the scrambled peptide had no further effect on staphylococcal adherence. Knockout of a different tetraspanin, TSPAN15, also reduced staphylococcal adherence to untreated cells (24.5±10.5%; Fig. S1C) but 800C was still inhibitory (49.4±6.4%; Fig. S2A). This peptide effect on adhesion is CD9-specific, as treatment of WT cells with a homologous peptide derived from tetraspanin CD81 had no effect (Fig. S3A, B). In addition, fluorescently-tagged 800C specifically bound to WT cells but not CD9^−/−^ cells (Fig. S3C-E). The inhibition of adhesion by 800C was also observed with a clinically relevant *S. aureus* strain, MRSA1 (Fig. S4A). Thus, we have identified a staphylococcal adherence pathway that can be inhibited by 800C only in the presence of CD9 on host cells.

**Figure 1.**
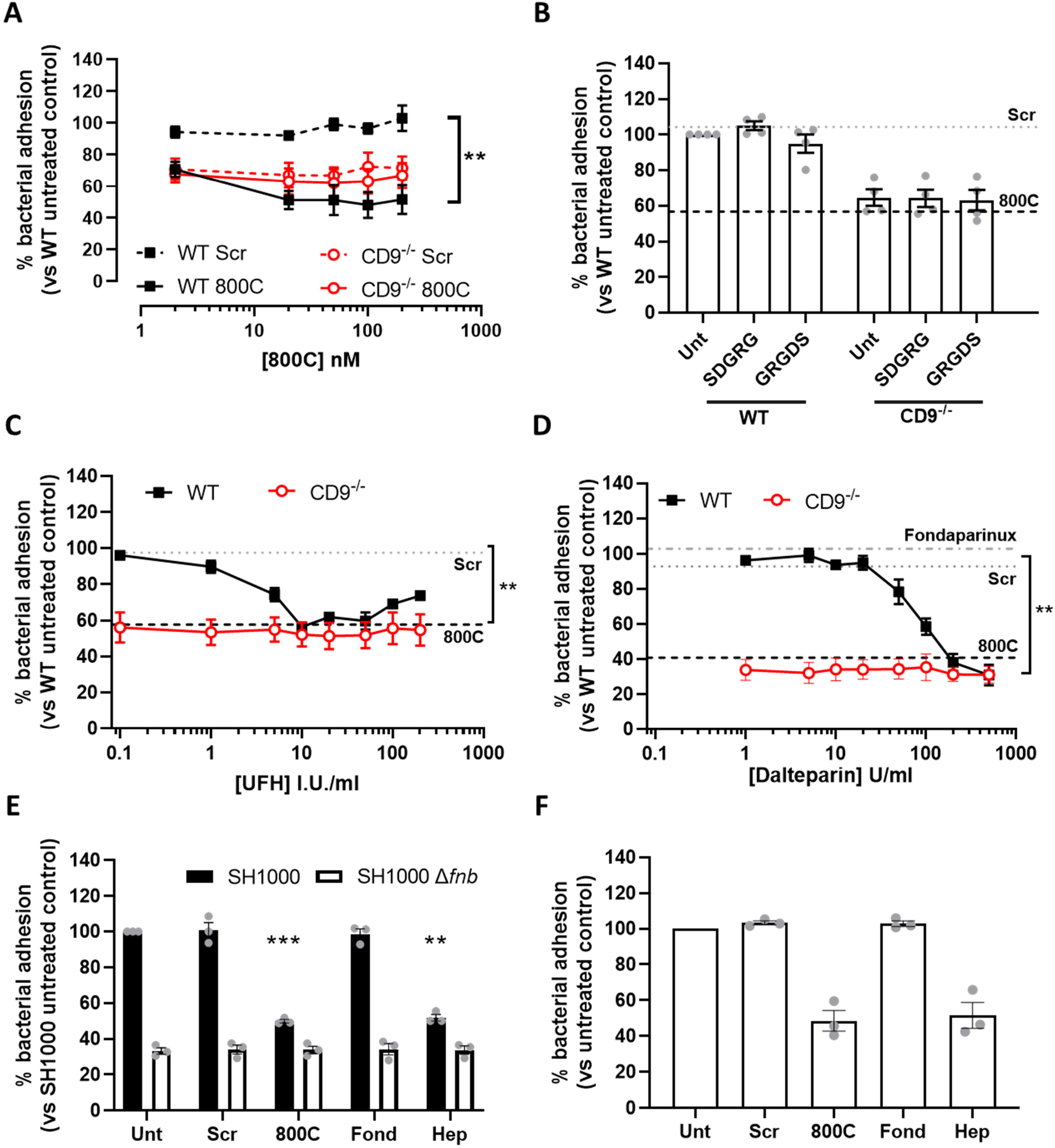
Tetraspanin derived peptides and heparin derivatives but not RGD peptides reduce staphylococcal adherence. Cells were infected with SH1000 for 60 mins at an MOI=50. (A) WT (black) or CD9^−/−^ (red) cells treated with scrambled (dotted) or 800C peptide (solid) for 60 mins prior to infection. WT (black) or CD9^−/−^ (red) cells treated with RGD peptides (100μM) (B), or various concentrations of heparin sodium (C) or dalteparin (D) for 60 mins prior to infection. Effect of CD9-derived peptide treatment (200nM) shown by dotted lines (B-D). Effect of fondaparinux (10μg/ml) treatment on WT cells shown in panel D. (E) WT cells treated with scrambled peptide, 800C, fondaparinux or UFH were infected with either SH1000 (black bars) or SH1000 Δ*fnb* (white bars) for 60 minutes at an MOI=50. (F) NTKs treated with scrambled peptide, 800C, fondaparinux or UFH were infected with SH1000 for 60 minutes at an MOI=50. n≥3, mean ± SEM, One-Way ANOVA.

### Staphylococcal adhesion is inhibited by heparins but not by integrin blockade

Despite previous studies demonstrating the importance of β1 integrins for staphylococcal invasion of epithelial cells, RGD peptides, which inhibit integrin-ligand interactions, had no effect on staphylococcal adhesion to epithelial cells (Fig. 1B). A similar lack of effect was observed with various anti-α5β1 integrin antibodies (P1D6, Lia1/2, JBS5 and AIIB2) (Fig. S5). In contrast, WT cells treated with unfractionated heparin (UFH) had significantly reduced staphylococcal adherence at 10 I.U./ml (43.5±2.2%), confirming a role for HSPG in staphylococcal adhesion. This reduction by UFH was biphasic, with maximal inhibition at 50 I.U./ml but less effect at higher concentrations. Similar results were observed for the keratinocyte cell line, HaCaT, with staphylococcal adhesion recovering almost completely at higher concentrations of UFH (Fig. S4B). Interestingly, no effect of UFH was observed with CD9^−/−^ cells (Fig. 1C), demonstrating a requirement for CD9 in UFH activity. Reductions in staphylococcal adherence to WT cells but not CD9^−/−^ cells were also observed when cells were treated with the low molecular weight heparin (LMWH), dalteparin (Fig. 1D). However, fondaparinux, a synthetic pentasaccharide based on the antithrombin binding region of heparin did not appear to affect adherence of *S. aureus* to WT or knockout cells (Fig. 1D). Similar effects of 800C and UFH were observed if the number of bacteria associated with epithelial cells were enumerated by microscopy (Fig. S4F). Infection with a *S. aureus Δfnb* deletion mutant, with both fibronectin binding protein A and B removed, demonstrated significantly reduced adherence to WT cells (Fig. 1E; 66.7±3.1%). Removal of these adhesins completely ablated the effects of both 800C and UFH (Fig. 1E). Similar effects of 800C and UFH were also observed with primary normal human tonsillar keratinocytes (NTK) that express high levels of CD9 and low levels of α5β1 and SDC-1 (Fig. 1F; Fig. S1D). Furthermore, 800C and UFH were also effective during infection of corneal and intestinal epithelial cell lines (Table S1), demonstrating the importance of both CD9 and HSPGs but not α5β1 integrin during staphylococcal adherence to a range of epithelial cell types.

### Heparin and CD9-derived peptides affect similar staphylococcal adherence pathways

The lack of effect of heparin and its analogues and 800C in CD9^−/−^ cells together with the observation that these treatments all reduce staphylococcal adherence to similar levels with WT cells (Fig. 1) suggests that these treatments affect similar pathways. We therefore tested 800C and the various heparin analogues in combination to test for additive effects. WT cells pre-treated with a combination of scrambled peptide and 10 I.U./ml UFH before infection demonstrated a significant reduction in staphylococcal adherence (54.7±19.4%); as expected these effects were smaller when treated with 5 I.U./ml of UFH. When UFH was combined with 800C, staphylococcal adherence was reduced to similar levels as either treatment alone (51.1±20.9%) with no evidence of a synergistic effect. Again, no effect of either treatment was observed with CD9^−/−^ cells (Fig. 2A). When WT cells were pre-treated with scrambled peptide and UFH was added during the infection, similar reductions in staphylococcal adherence were observed when treated with UFH or 800C alone (52.2±7.5%; Fig. 2B). A combined treatment of UFH and 800C on WT cells caused no further additional effects while treatment of CD9^−/−^ cells had no effect. Similarly, pre-treatment of cells with LMWH dalteparin and scrambled peptide reduced staphylococcal adherence to similar levels as cells pre-treated with 800C or dalteparin alone (54.0±12.5%; Fig. 2C). Combination treatment of dalteparin and 800C had no further additive effect and treatments had no effect on CD9^−/−^ cells. Pre-treatment of cells with scrambled peptide and fondaparinux demonstrated no significant reductions in staphylococcal adherence and, when combined with 800C, adherence was reduced to levels similar to that of 800C treatment alone (54.2±14.9%; Fig. 2D). No effects of either treatment were observed with CD9^−/−^ cells or when WT cells were infected with a Δ*fnb* mutant (Fig. 2E). Similarly, no additive effects were observed after a combined treatment of UFH and 800C on NTKs (Fig. 2F). Thus, CD9 has been implicated within the same staphylococcal adherence pathway as the HSPGs in epithelial cells for the first time.

**Figure 2.**
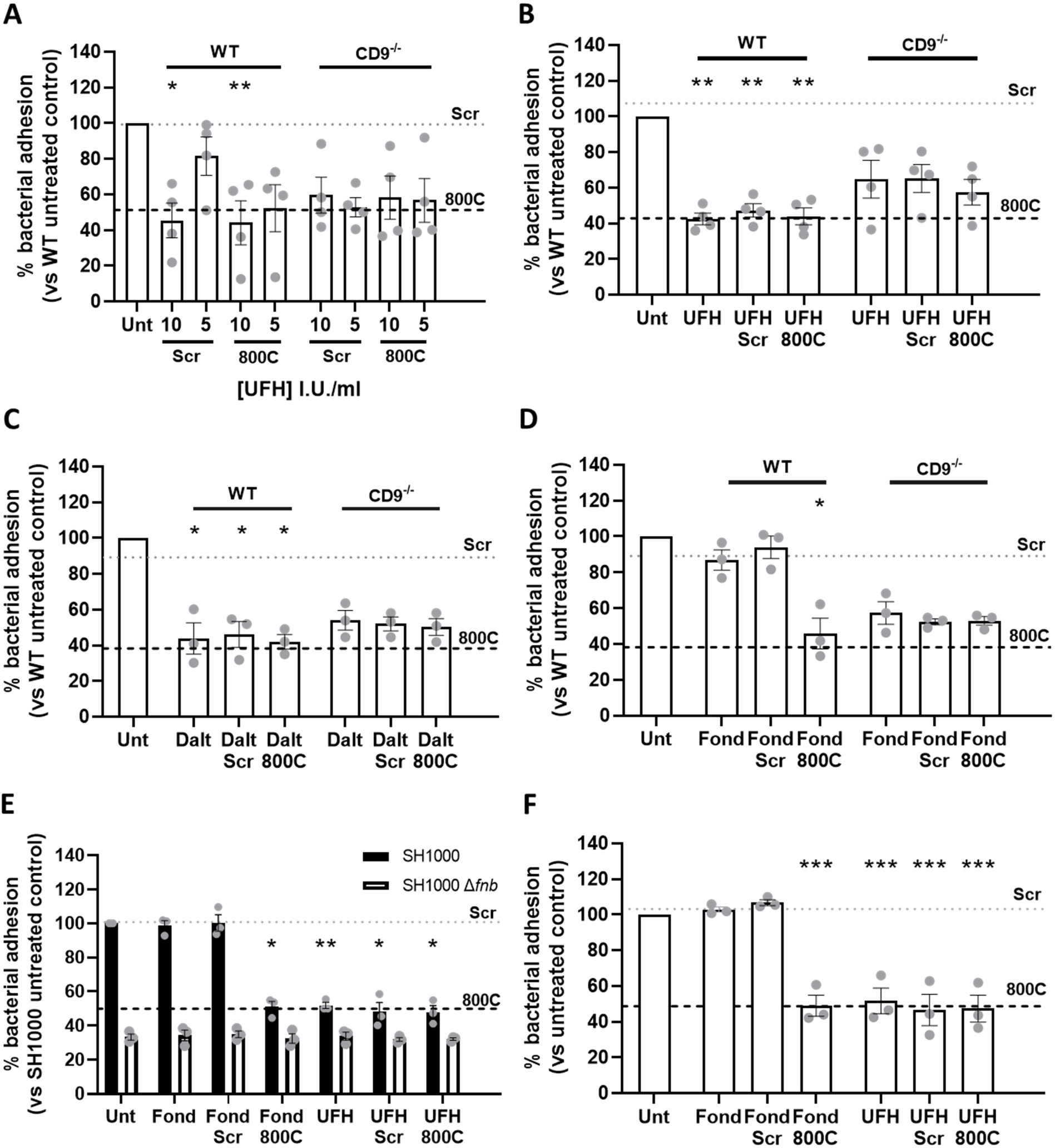
Combination treatment of unfractionated heparin and 800C produces no additive effects. Cells were infected with SH1000 for 60 mins at an MOI=50. (A) WT or CD9^−/−^ cells were treated with UFH, 800C peptide or a combination of both for 60 mins prior to treatment. (B) WT or CD9^−/−^ cells were treated with peptide (200nM) for 60 mins prior to infection. UFH (10 U/ml), either in combination with 800C peptide or as a singular treatment, was added at the start of infection. (C) WT or CD9^−/−^ cells were treated with dalteparin (200U/ml), 800C peptide (200nM) or a combination of both 60 minutes prior to infection. (D) WT or CD9^−/−^ cells were treated with fondaparinux (10μg/ml), 800C peptide (200nM) or a combination of both 60 minutes prior to infection. (E) WT cells treated with 800C peptide (200nM), UFH (10U/ml) or a combination of the two and infected with either SH1000 or SH1000 Δ*fnb* at an MOI=50. (F) NTKs treated with 800C peptide (200nM), UFH (10I.U./ml) or a combination of the two and infected with SH1000 for 60 minutes at an MOI=50. n≥3, mean ± SEM, One-Way ANOVA.

### Heparan sulphates are required during tetraspanin-mediated staphylococcal adherence

Reductions in staphylococcal adherence by both 800C and heparin analogues suggests that HSPGs are involved during tetraspanin-mediated staphylococcal adherence. To further test this, adherence of *S. aureus* to epithelial cells was measured after removal of HS from the cell surface using heparinases. Treatment with a mixture of heparinase I/III significantly reduced staphylococcal adherence to WT cells (53.1±13.7%), while combination treatment with scrambled peptide or 800C had no further additive effect (Fig. 3A). Treatment of WT cells with chondroitinase ABC did not reduce staphylococcal adherence. No significant reduction was observed when CD9^−/−^ cells were treated with heparinase or chondroitinase (Fig. 3B). Staphylococcal adherence was also significantly reduced when cells were pre-treated with 10 μg/ml HS (37.3±4.2%; Fig. 3C) or DS (42.1±3.4%; Fig. 3E). As with heparin, treatment of CD9^−/−^ cells with HS or DS did not reduce staphylococcal adherence. CS treatment had no effect on WT or CD9^−/−^ cells (Fig. 3D). Treatment with pixatimod, a clinical stage HS mimetic (Dredge et al., 2018), also caused a significant reduction in staphylococcal adhesion to WT cells (42.4±4.5%; Fig. 3F) but not CD9^−/−^ cells. This suggests the importance of HS chains but not the core protein of HSPGs, during CD9-mediated staphylococcal adherence, and demonstrates a potential new therapeutic to interfere with this process.

**Figure 3.**
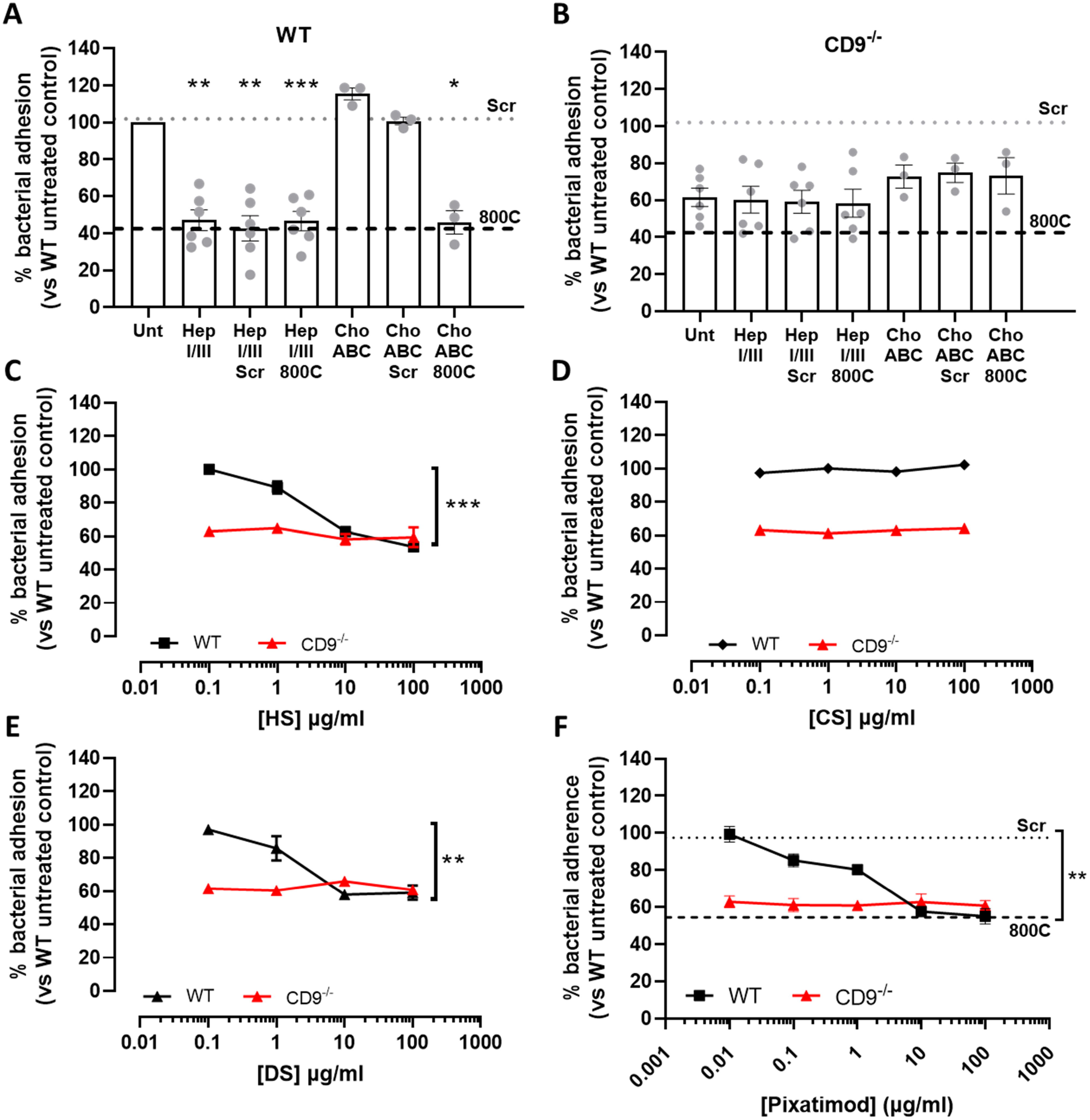
Heparan sulphates are required during tetraspanin-mediated staphylococcal adherence. Cells were infected with SH1000 for 60 mins at an MOI=50. (A) WT cells were treated with either 0.25U/ml chondroitinase ABC or 0.5U/ml heparinase I/III for 3 hours prior to infection. Peptide was added to cells 60 mins prior to infection for combination treatments. (B) CD9^−/−^ cells were treated with either 0.25U/ml chondroitinase ABC or 0.5U/ml heparinase I/III for 3 hours prior to infection. Peptide was added to cells 60 mins prior to infection for combination treatments. WT or CD9−/− cells were treated with heparan sulphate (HS;C), chondroitin sulphate (CS; D), dermatan sulphate (DS; E) or the heparan sulphate mimetic, pixatimod, (F) for 60 minutes prior to infection. n≥3, mean ± SEM, One-Way ANOVA.

### Tetraspanin-mediated staphylococcal adherence requires SDC-1

Removal of HS from HSPG, treatment with heparin analogues or with HS all reduce staphylococcal adherence in a similar manner to 800C, suggesting the involvement of HSPGs in tetraspanin-mediated adherence. A number of HSPGs have been demonstrated to associate with CD9, including SDC-1 (Jones et al., 1996). Syndecans are expressed on epithelial cells, with SDC-1 and SDC-4 the most abundant (Kim et al., 1994). Here, we tested the involvement of SDC-1 and −4 in this process using blocking antibodies and shRNA knockdowns. Both SDC-1 and −4 were expressed on WT cells, with SDC-1 expression higher than SDC-4 (Fig. S1A). Staphylococcal adherence was reduced after treatment with an anti-SDC-1 antibody (46.6±3.3%) similar to treatment with 800C (Fig. 4A) but no effect was observed if WT cells were treated with an isotype control antibody. No further additive effect was observed if the anti-SDC-1 antibody was added in combination with 800C, while antibody treatment of CD9^−/−^ cells had no effect. Anti-SDC-4 antibodies had no effect on staphylococcal adherence (Fig. 4B). Partial knockdown of either SDC-1 or SDC-4 was very efficient (Fig. S6) but had only small inhibitory effects on staphylococcal adherence to epithelial cells (29.6±9.4% and 19.0±10.1%, respectively) (Fig. S1C), suggesting some redundancy between the SDC. However, knockdown of SDC-1 negated the effects of 800C and UFH on WT cells relative to untreated or scrambled shRNA treated cells (Fig. 4C) whereas 800C and UFH were still effective at reducing staphylococcal adherence to SDC-4 knockdown cells (77.1±6.0% and 78.7±7.4%, respectively). Thus, SDC-1, a known partner protein of CD9 (Jones et al., 1996), appears to be the primary membrane protein required for CD9-mediated staphylococcal adhesion.

**Figure 4.**
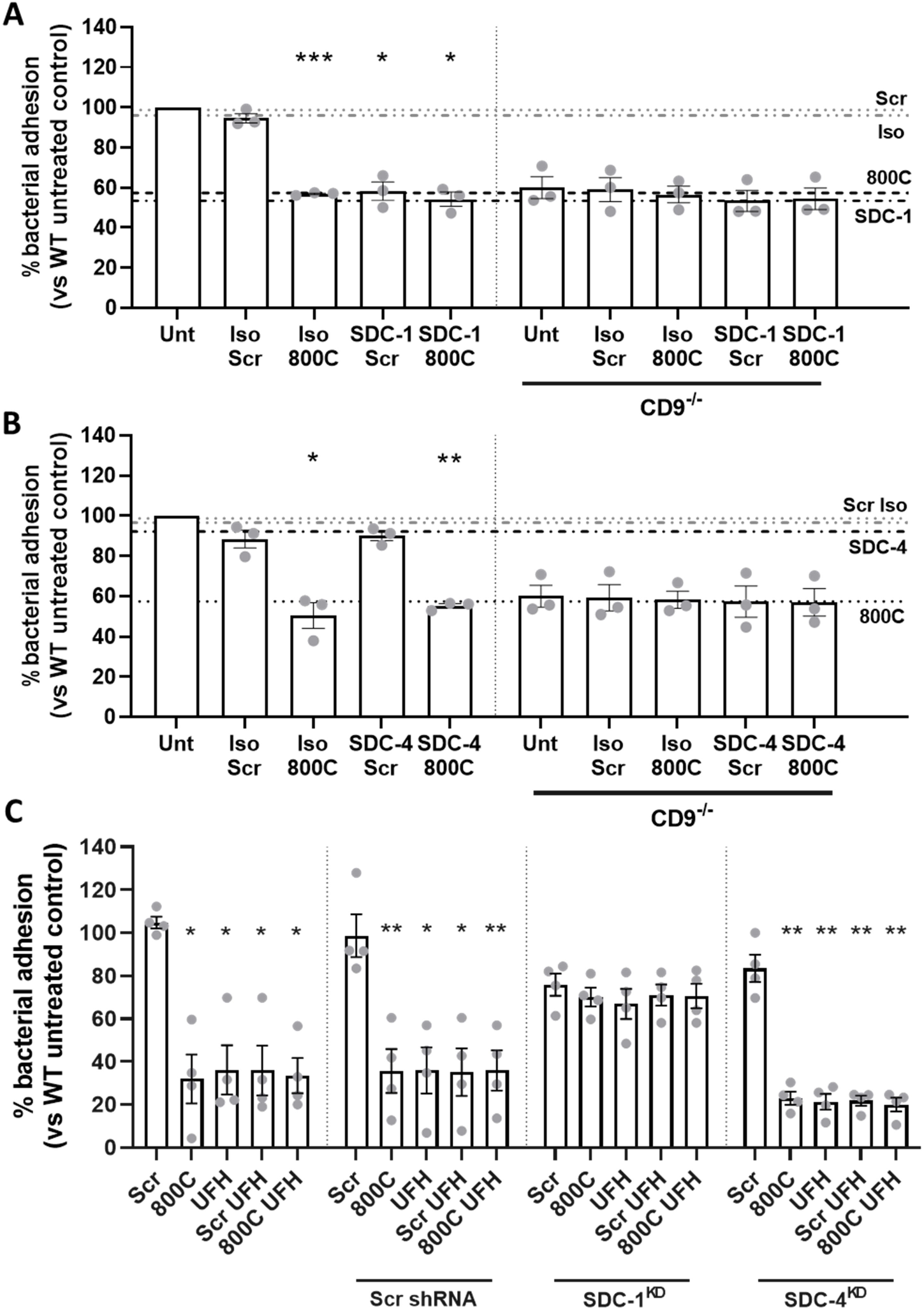
Syndecan-1 is involved in tetraspanin-mediated staphylococcal adherence to epithelial cells. Cells were infected for 60 mins with SH1000 at an MOI=50. (A) WT or CD9^−/−^ cells were treated with peptides (200nM), isotype control (JC1), anti-SDC-1 antibodies (B-A38, 10μg/ml) or a combination of peptide and antibodies for 60 mins prior to infection. (B) WT or CD9^−/−^ cells were treated with peptides (200nM), isotype control (02-6200), anti-SDC-4 antibodies (5G9, 20μg/ml) or a combination of peptide and antibodies for 60 mins prior to infection. (C) shRNA knockdowns of SDC-1 and SDC-4 were treated with peptides (200nM), UFH (10I.U./ml) or a combination of the two for 60 mins prior to infection. Scrambled shRNA was used as a control. n≥3, mean ± SEM, One Way ANOVA.

### Interference with HSPGs and CD9-derived peptide increases staphylococcal internalization

We have demonstrated that staphylococcal adherence can be reduced by interfering with HSPGs and CD9 but not by blockade of β1 integrins. Here, we tested the effects of these treatments on the internalization of *S. aureus* into epithelial cells using a gentamicin protection assay. For CD9^−/−^ cells, internalization of adherent bacteria was significantly increased (2.7±0.6%; *p*=0.0001) compared to WT cells (1.4±0.34%; Fig. 5A). A similar increase was observed with WT cells treated with 800C (2.0±0.6%); however, this did not reach significance (p=0.094). This increase could be due to changes in the number of cell-associated bacteria but no increase was observed with Tspan15^−/−^ cells (Fig. 5A), where reductions in cell-associated bacteria were also observed.

**Figure 5.**
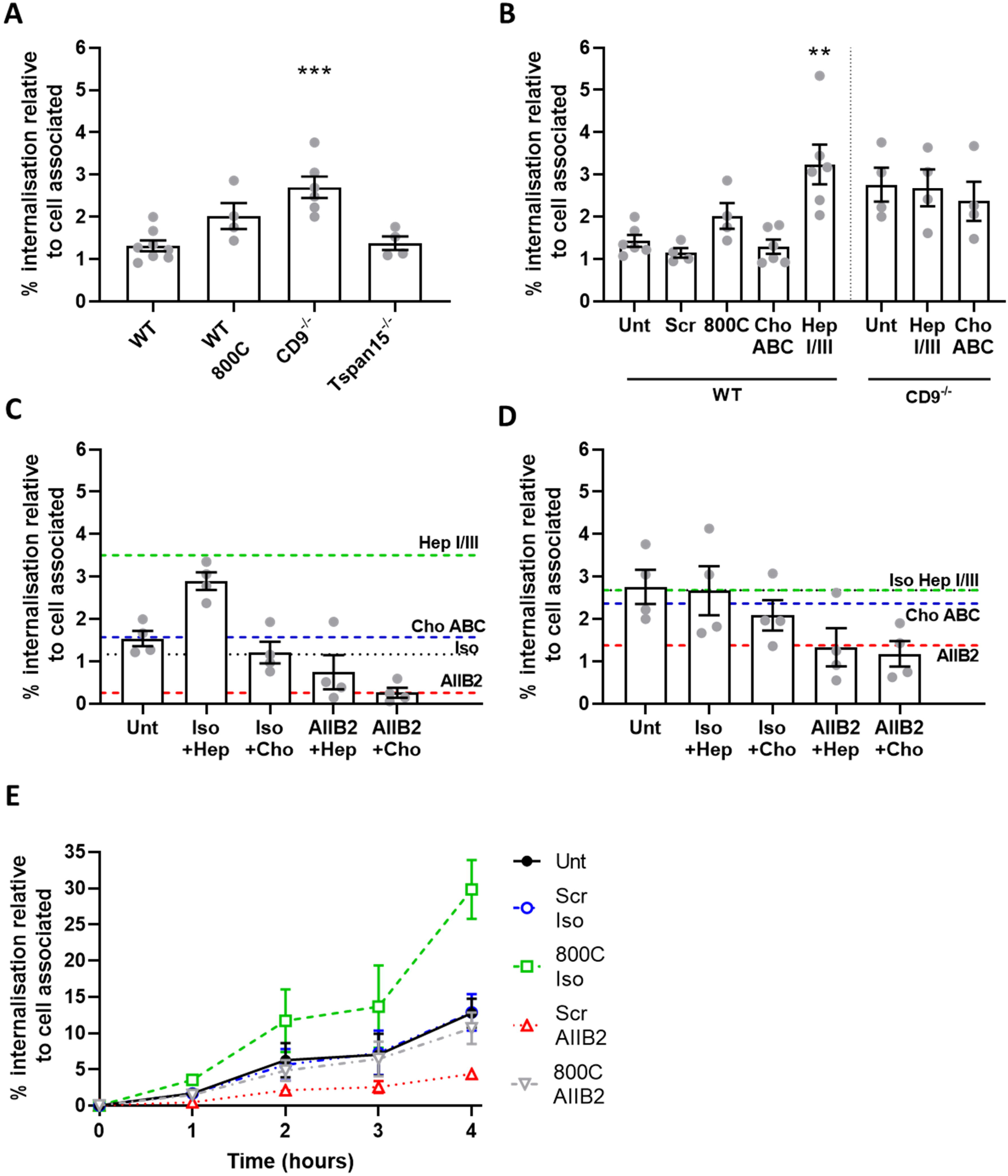
Interference with CD9 or heparan sulphates increases staphylococcal internalisation, which is controlled by α5β1. Cells were infected with SH1000 for 60 mins at an MOI=50. After infection, cells were washed and treated with 200μg/ml gentamicin sulphate to remove adherent bacteria and lysed to quantify the internalised bacteria. Internalised bacteria were normalised against the cell associated bacteria. (A) Internalised bacteria within WT, CD9^−/−^ and Tspan15^−/−^ cells, 800C treated WT cells are added as an example. (B) WT or CD9^−/−^ cells were treated with 0.25U/ml chondroitinase ABC or 0.5U/ml heparinase I/III for 3 hours prior to infection. WT (C) or CD9^−/−^ (D) cells were treated with heparinase I/III or chondroitinase ABC for 3 hours. Cells were treated with isotype control or anti-α5β1 antibodies (AIIB2) for 60 minutes prior to infection. (E) Cells were treated with combinations of peptide (200nM), isotype control, anti-α5β1 antibodies (AIIB2) for 60 minutes prior to infection. Cells were infected for 60 mins at 4°C, after infection cells were warmed to 37°C and internalisation was allowed to continue for 4 hrs. n≥3, mean ± SEM, One-Way ANOVA.

Increases in internalization were also observed in WT cells treated with heparinase I/III (3.2±1.2%) but not with chondroitinase ABC (Fig. 5B), whereas no further increases were observed for CD9^−/−^ cells treated with heparinase I/III. Treatment of WT cells with AIIB2, an antibody able to dissociate β1:Fn complexes, reduced internalization to negligible levels (0.3±0.2%) while treatment of CD9^−/−^ cells with AIIB2 only reduced internalization to levels observed in untreated WT cells (Fig. 5C,D). Increases in internalization were observed for 4 hours after infection (Fig. 5E), although increasing cell death prevented measurement after this point. Treatment of WT cells with a combination of a scrambled peptide with AIIB2 reduced internalization to very low levels throughout, similar to the AIIB2 alone. Interestingly, treatment of WT cells with a combination of 800C and AIIB2 reduced tetraspanin-mediated internalization back to levels of internalization observed with untreated WT cells at all time points (Fig. 5E), suggesting that β1:Fn complexes are still critical for internalization of *S. aureus* even in the absence of CD9. Similarly, if AIIB2 is used in combination with heparinase treatments, internalization is reduced to untreated levels (0.8±0.8%; Fig. 5D). These data indicate that CD9 is a positive regulator of HSPGs during initial adherence but that CD9 is a negative regulator of the subsequent α5β1-mediated internalization, suggesting that CD9 links these two separate events.

### Changes in the expression of SDCs and α5βl integrin does not explain the reduced staphylococcal adherence in CD9 knockout cells

The expression of α5β1 integrins and the SDCs was measured by flow cytometry in CD9^−/−^ cells to determine if the effects on staphylococcal adherence is secondary to changes in expression of these membrane proteins. As expected for CD9^−/−^ cells, CD9 expression was dramatically reduced compared to WT cells (−73.3±1.5%; Fig. 6A) with the number of positive cells reduced by 95.4±0.5% (Fig. S1B). Surprisingly, both SDC-1 and SDC-4 demonstrated increased expression on CD9^−/−^ cells (117.5±25.8% and 21.6±11.3% respectively). although the number of SDC-4 positive cells dropped markedly (50.8±5.8%). Integrin α5 and β1 expression was measured with two separate antibodies; AIIB2 is able to dissociate β1:Fn interactions and can therefore measure total surface β1, and JBS5, which is unable to dissociate these interactions and so only measures the unliganded fraction of α5 (Mould et al., 2016). Interestingly, JBS5 binding was reduced in CD9^−/−^ cells relative to WT (−44.9±15.0%; Fig. 6A) with the percentage of positive cells also reduced (−89.9±1.8%; Fig. S1B) whilst AIIB2 binding was increased in the CD9^−/−^ cells (9.9±9.0%). These data suggest that CD9 can regulate cell surface expression of SDCs but SDC expression levels are not the critical determinant of staphylococcal adherence. CD9 may also act to keep a population of α5β1 in an inactive state unable to bind Fn.

**Figure 6.**
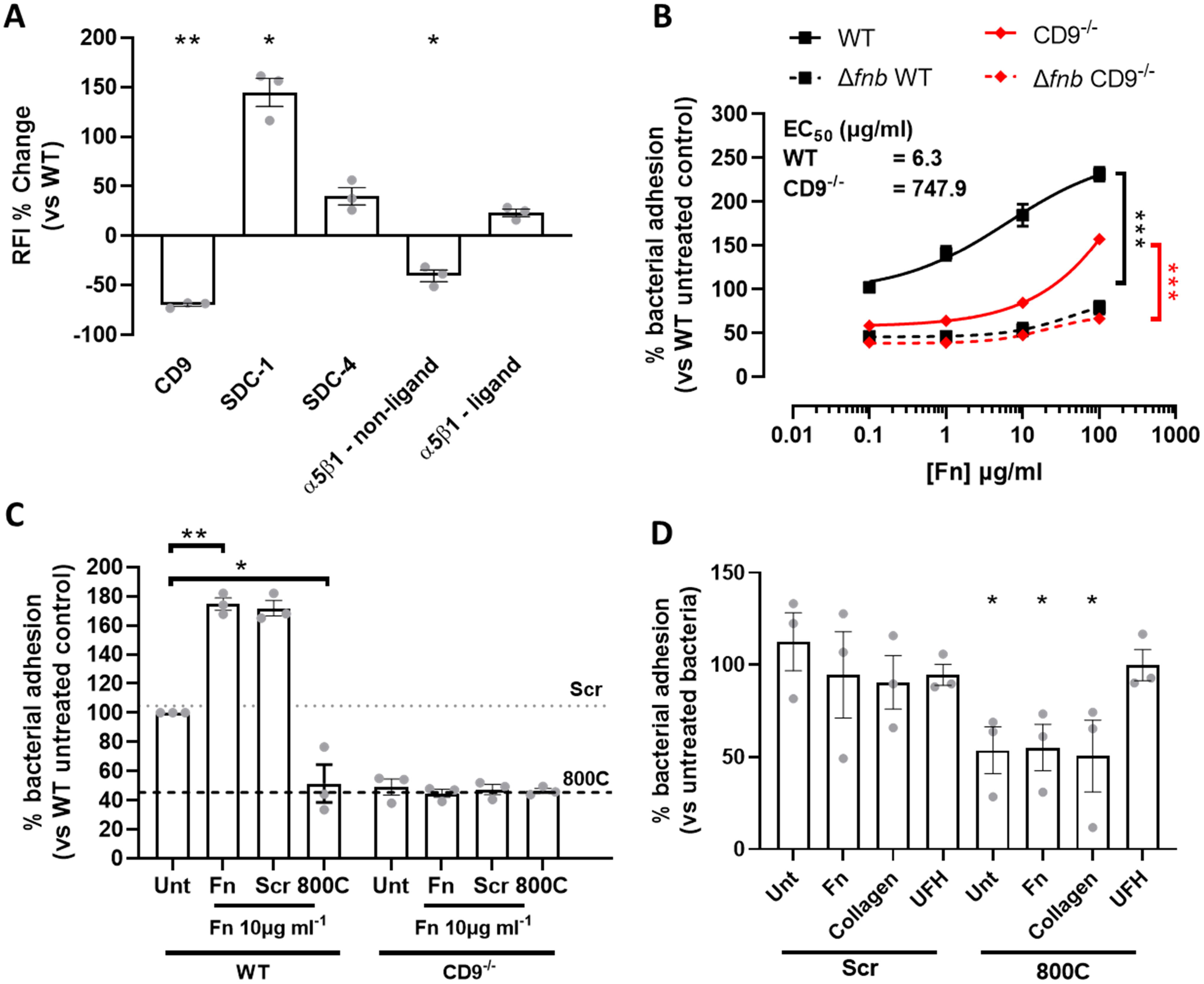
Addition of exogenous Fn increases staphylococcal adherence, which is negated in the presence of 800C. (A) Cell surface expression of receptors of interest was determined in CD9^−/−^ cells. Relative fluorescence intensity was calculated by dividing the test antibody by the isotype control. Percentage change was calculated against WT cell values. (B) Cells were treated with varying concentrations of Fn prior to infection with SH1000 or SH1000 Δ*fnb*. (C) Fn was added to WT or CD9^−/−^ cells in combination with 800C or a scrambled peptide for one hour prior to infection. (D) SH1000 was pre-treated with Fn, collagen or UFH for 90 minutes prior to infection. Cells were treated with 200nM 800C or scrambled peptide for 60 minutes prior to infection. n≥3, mean ± SEM, One-Way ANOVA.

### Fn is important for tetraspanin-mediated staphylococcal adherence

As differences were observed in cell surface proteins known to bind extracellular matrix proteins we tested whether bacterial adherence could be affected by the addition of exogenous Fn. The addition of Fn to WT cells increased staphylococcal adherence in a dose-dependent manner, with only low concentrations of Fn required to increase *S. aureus* binding to WT cells (EC_50_ = 6.3 μg/ml, Fig. 6B). However, a much higher concentration of Fn is required to increase staphylococcal adherence to CD9^−/−^ cells (EC_50_= 747.9 μg/ml). Adherence by the Δ*fnb* mutant remained unaffected by the addition of exogenous fibronectin. These data are not explained by increases in non-specific binding to plastic wells treated with Fn (Fig. S7). To further test the importance of CD9 during increased binding of *S. aureus* after addition of exogenous extracellular matrix proteins, we tested 800C in combination with 10 μg/ml Fn, which reduced staphylococcal adherence (51.5±22.4%) to levels similar to that of 800C treatment alone, despite the increases observed if Fn was added alone (175±7.20%) or in combination with the scrambled peptide (172α9.10%; Fig. 6C). No effect of Fn or 800C was observed with CD9^−/−^ cells (Fig. 6C) or after infection with a Δ*fnb* mutant (Fig. S7B). Interestingly, addition of UFH, Fn or collagen to bacteria prior to infection had no effect on staphylococcal adherence to cells (Fig. 6D) although adherence of Fn or collagen-treated bacteria was significantly reduced after treatment with 800C (44.8±21.8% and 49.4±33.7%, respectively). Interestingly, the effect of 800C was completely abrogated when cells were infected with UFH treated bacteria. Taken together, this suggests a significant role for Fn during CD9-mediated staphylococcal adherence.

## Discussion

In this study, we have identified a critical role for CD9 in the control of the behaviour of SDC-1-dependent *S. aureus* adhesion sites. Integrin α5β1 plays no part in this adherence pathway but does have a role in the subsequent internalization of bacteria. Interference with CD9/SDC-1 led to increases in internalization that were reduced by integrin blockade, suggesting a sequential CD9-mediated process connecting adherence with internalization. The addition of exogenous Fn increased binding in a CD9-sensitive manner, suggesting the importance of Fn during tetraspanin-mediated adherence, likely by integration with CD9/SDC-1. Finally, we present two potential anti-adhesive therapeutics, CD9-derived peptide 800C and heparin, providing a mode of action for both.

### A new model for staphylococcal adhesion to epithelial cells

As previous studies have demonstrated direct interaction of CD9 with either α5β1 or SDC-1 by co-immunoprecipitation (Jones et al., 1996) we propose the following model (Fig. 7), in which CD9 TEMs contain either SDC-1 or α5β1. Clustering of SDC-1, within CD9 enriched microdomains, leads to recruitment and organisation of Fn fibrils into ‘adhesion nets’ on the cell surface (Stepp et al., 2010; Jinno et al., 2020) increasing bacterial adherence. CD9-blockading reagents, such as 800C and antibodies, may operate by disrupting CD9-containing TEMs and therefore the organisation of Fn fibrils by TEM-associated SDC-1. Treatment with UFH or heparin derivatives may also disrupt the established fibronectin adhesion net (Galante and Schwarzbauer, 2007) or block staphylococcal association to Fn (Arciola et al., 2003) therefore reducing bacterial adhesion efficiency. 800C and UFH treatments were both able to reduce staphylococcal adherence by approximately 50-60%, demonstrating that other adherence pathways remain unaffected by these treatments.

**Figure 7.**
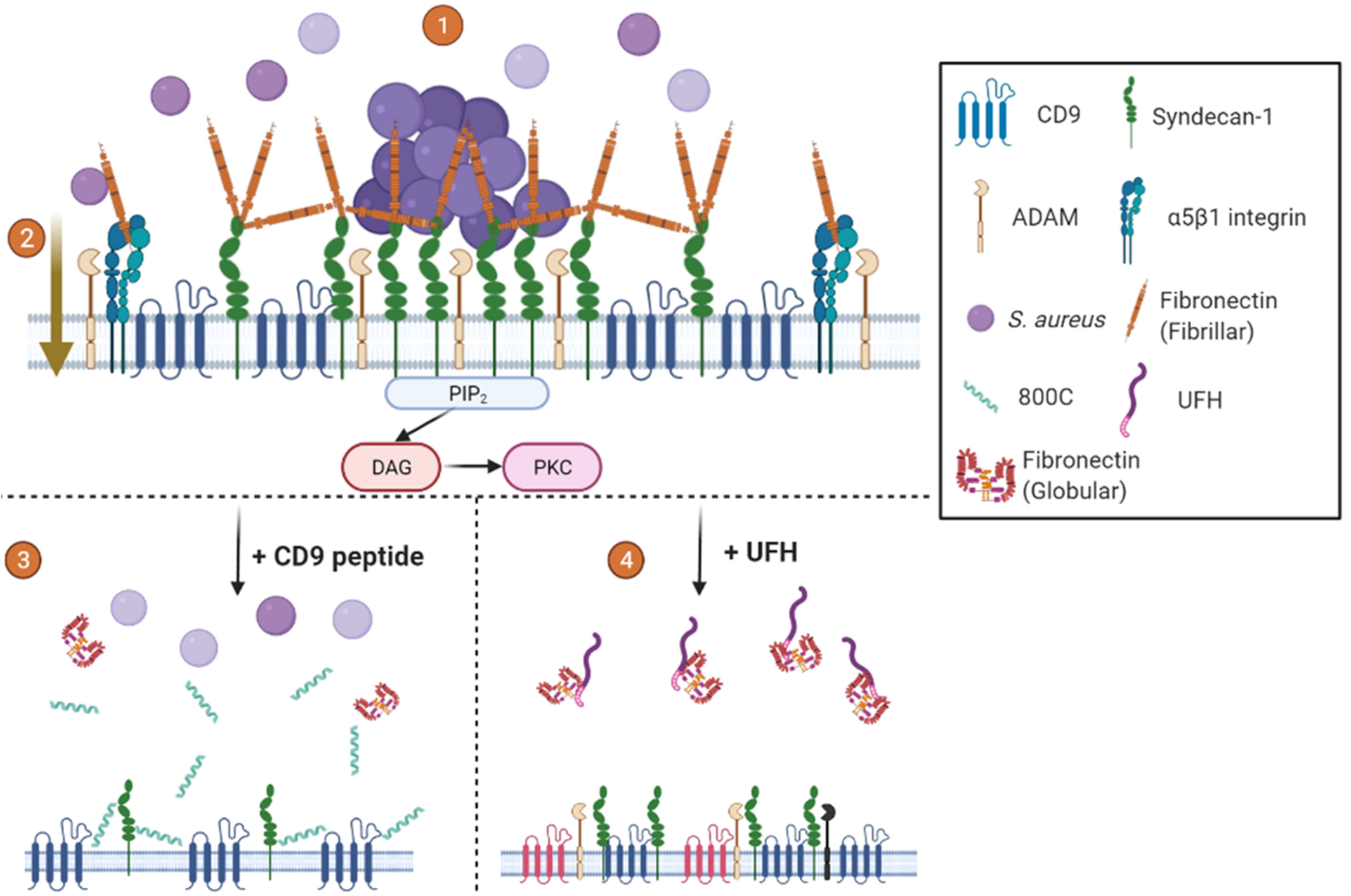
Proposed mechanism for tetraspanin-mediated staphylococcal adherence. (1) CD9 enriched microdomains contain syndecan-1. Clustering of syndecan-1 by CD9 recruits fibronectin and produces ‘adhesion nets’. *S. aureus* utilises these nets to initiate initial adherence to the cell surface. (2) Inclusion of integrins in CD9 enriched microdomains means the ‘adhesion nets’ are in close proximity to β1 integrins, allowing for transfer of bacteria after initial adherence for rapid internalisation through the canonical pathway. Interactions with CD9 and ADAMs ensures integrins are kept in an inactive state inhibiting bacterial internalisation. (3) CD9-derived peptides increase the area of TEMs pushing syndecan clusters apart and reducing formation of ‘adhesion nets’. (4) UFH treatment displaces fibronectin or blocks staphylococcal interaction with Fn, destabilising TEMs and reducing bacterial adhesion. Mechanisms other than CD9-mediated adhesion and internalisation are still possible with both treatments. Created with Biorender.com.

### The role of syndecans in staphylococcal adhesion

Our findings add to others that have implicated HSPGs, and particularly SDC-1, in the staphylococcal adherence pathway (Hess et al., 2006; García et al., 2016; Rajas et al., 2017). However, some studies have suggested that while SDC-1 is important for staphylococcal infection, it is not a direct receptor. For example, Hayashida *et al.* demonstrated a lower burden of staphylococcal corneal infection in Sdc1^−/−^ knockout mice but did not observe interaction of *S. aureus* with SDC-1 on beads or with a mouse SDC-1 knockdown corneal epithelial cell line (Hayashida et al., 2011). They instead suggest that *S. aureus* induction of SDC-1 ectodomain shedding, likely through the action of ADAMs (a disintegrin and metalloproteinase) or other metalloproteinases, is critical for improved survival of the bacteria. SDC-1 ectodomains have been shown to inactivate cathelicidins and reduce neutrophil-mediated killing, significantly increasing staphylococcal survival during infection (Park et al., 2001; Hayashida et al., 2015). The association of tetraspanins with syndecans and ADAMs suggest the tetraspanins may also be involved in ectodomain shedding. Further studies investigating the role of CD9 in ADAM-mediated ectodomain shedding could provide valuable data into tetraspanin control of another putative pathogenesis pathway. However, we would further suggest the expression profile between the cell lines used and those in this study differ significantly with lower levels of CD9 potentially reducing clustering of receptors and so lowering the avidity of interactions between *S. aureus* and SDC-1. This study has utilised multiple cell lines which show both differences in staphylococcal adhesion and protein expression profile (Table S1). Furthermore, whilst we have shown that SDC-1 appears to be important during tetraspanin-mediated adhesion, we also observed an increase in SDC-1 expression on CD9^−/−^ cells (Fig. 6A), to which staphylococcal adhesion is also significantly reduced suggesting that coordination and clustering of this protein is important.

Other studies suggest that the primary role of SDC-1 is the organisation of Fn at the host cell surface, through the GAG binding domains of Fn (Jinno et al., 2020). Various bacteria are then able to interact directly with Fn through FnBPs located in their outer membrane (Speziale et al., 2019). Our data suggest that the coordinate action of CD9 and SDC-1 is required for efficient staphylococcal adherence, whereby CD9 clusters SDC-1 in TEM to allow the organisation of Fn into adhesion platforms. However, we cannot rule out that a specific moiety of HS is required on SDC-1 for adherence, as we demonstrate that HS as well as DS, which both decorate syndecans, reduce staphylococcal adherence to a similar extent as 800C. Whilst we have not investigated other HSPGs such as the glypicans. These GPI-anchored proteins are usually present in lipid rafts (Mayor and Riezman, 2004), unlike the syndecans that have been associated with TEMs (Jones et al., 1996; Grigorov et al., 2017), and so glypicans are less likely to be involved in these processes. Furthermore, the equivalent effects of heparins and the CD9-derived peptide, together with the abrogation of these effects on SDC-1 knockdown suggests that the glypicans cannot replace SDC in this tetraspanin-mediated adherence pathway.

### The role of integrins in staphylococcal adhesion and internalization

Syndecans and integrins interact with Fn and other extracellular matrix proteins to provide structural support enabling cell:cell adhesion and transmembrane interactions for downstream cell signalling (Couchman, 2010). Both families of proteins have been implicated in Fn fibril assembly, a process thought to require co-localisation and clustering of these proteins (Morgan et al., 2007; Araki et al., 2009; Stepp et al., 2010). Loss of SDC-1 in corneal cells led to less activated β1 integrin, which was used to explain reduced Fn fibril formation (Stepp et al., 2010). In addition, expression of CD9 on Chinese hamster ovary cells led to reduced cell adhesion and increased spreading on Fn (Cook et al., 1999), with CD9 expression proposed to stabilise the ‘active’ conformation of α5β1 (Kotha et al., 2008). Interestingly, Cook et al found that a 37 aa peptide derived from the CD9 EC2, containing the sequence utilised in the present study, was able to reverse the inhibition of CHO cell adhesion to Fn (Cook et al., 2002). However, our data suggest that CD9-mediated SDC clustering may provide an indirect link between CD9 and Fn. CD9 has also been shown to negatively regulate cell adhesion by promoting the interaction of α5β1 with ADAM17, keeping both molecules in an inactive state (Machado-Pineda et al., 2018). No change in the activation state of α5β1 was observed but α5β1 shifted from an equal distribution pattern on the cell surface to more punctate clusters in the absence of CD9 (Machado-Pineda et al., 2018). The FnBPs of *S. aureus* are known to drive the clustering of integrins upon interaction with Fn, which leads to rapid internalization (Liang et al., 2016). The present study focuses on changes within host adhesion receptors but poses intriguing questions as to the role of *fnbA* and *fnbB* within tetraspanin-mediated staphylococcal adhesion. Here, we showed that ligand-bound β1, suggesting an ‘active’ integrin conformation, increased in the absence of CD9; however, this was accompanied by higher expression of SDC-1. We also observed greater staphylococcal adherence in the presence of Fn, which was abrogated by the CD9-derived peptide, and increased internalization with the loss of CD9 or after interference with HSPGs. Recently and in agreement with our studies, the supramolecular structure of fibronectin at the cell surface has been linked to differential uptake of *S. aureus* (Niemann et al., 2021). A denser, layered Fn fibril network at the surface of osteoblasts demonstrated a poorer uptake of bacteria compared to a moderate Fn fibril network at the surface of A549 epithelial cells suggesting an optimal concentration and organisation of Fn is required for efficient adhesion and uptake. We therefore suggest that clustering of SDC-1 by CD9 and the resulting Fn fibril formation drives the initial adherence of *S. aureus.* These bound bacteria are then able, through the action of FnBPs, to begin clustering of α5β1 and thus promote internalization over adherence.

### The potential of CD9 peptides and heparins as anti-infective agents

We have provided insights into the mode of action of a new anti-bacterial adhesive therapeutic, 800C, suggesting that it may disrupt SDC-1 clusters that act as the anchor of ‘adhesion nets’ on the cell surface, significantly inhibiting staphylococcal adherence to a variety of host cell types. Future studies will investigate analogues of 800C to further reduce staphylococcal adherence and to check safety and efficacy of tetraspanin-derived peptide treatments. We have recently demonstrated that a stapled form of 800C is effective in *in vivo* and *ex vivo* models of infection with no detrimental effects to the host (Patent: WO2021/175809A1). UFH and heparin analogues have previously been demonstrated to inhibit the adherence of various bacteria and viruses (Aquino and Park, 2016). Our study adds significantly to the growing body of evidence that suggests heparin could be used as an anti-adhesive therapeutic and provides a mechanism for its mode of action. Previously thought to interact directly with some bacterial adhesins (Chen et al., 2008) or with fibronectin (Arciola et al., 2003), our data suggests that heparin also acts on the host via the SDC/CD9/Fn ‘adhesion net’. Whilst we observed a biphasic response of UFH upon staphylococcal adherence suggesting that heparin may be directly utilised by the bacteria at higher concentrations (Shanks et al., 2005), we also demonstrated inhibitory effects of low molecular weight heparins. However, typical therapeutic serum concentrations of LMWH are 0.5-1.2 U/ml (Jaffer and Weitz, 2018), much lower than the inhibitory concentrations required in this study (Dalteparin IC50 = 85.2 U/ml). Fondaparinux, a synthetic pentasaccharide heparinoid able to bind antithrombin but not thrombin (Bauer et al., 2002), showed little effect here, providing an insight into the structural requirements for a heparin-based anti-infective. We have also demonstrated for the first time significant inhibition of staphylococcal adherence using Pixatimod (PG545, IC_50_ = 1.11 μg/ml), a clinical stage HS-mimetic that shows potent anti-cancer and anti-inflammatory effects (Hammond and Dredge, 2020). Pixatimod has also been reported to have significant anti-viral activity against a range of viruses which utilise HS as receptors including SARS-COV-2, HSV-2, HIV, RSV and Dengue (Said et al., 2010, 2016; Lundin et al., 2012; Modhiran et al., 2019; Guimond et al., 2022). The emergence of new HS mimetics with significantly reduced anticoagulant activity suggests new avenues of research to develop anti-adhesive therapeutics against existing and emerging pathogens.

In summary, we have made the novel observation that the presence of CD9 at the cell surface, alongside SDC-1 and the integrins, is a critical component of a major mechanism of *S. aureus* adherence to epithelial cells. We have also demonstrated the importance of Fn for CD9/SDC-1-mediated staphylococcal adherence and present two potential therapeutic pathways to significantly reduce staphylococcal adherence, namely CD9-derived peptides and heparin/HS analogues. With this study providing mechanistic details as to the action of these putative therapeutics, future studies should now investigate analogues which could further reduce staphylococcal adherence beyond the levels of inhibition demonstrated here. Previous data has suggested tetraspanin involvement in the adherence of a wide variety of Gram-positive and -negative bacterial pathogens and so further investigations may shed light on a common pathway for bacterial pathogenicity involving the organisation of HSPGs by tetraspanins. Furthermore, the tetraspanins are associated with and organise a number of different partner proteins that may act as bacterial host receptors; continued research may reveal more potential therapeutics relieving the burden during the antimicrobial resistance crisis.

## Author Contributions

Conceptualization, PNM, LJP, BC and LRG; Methodology, LRG, RI, FA, HK, CET, LJP and PNM; Formal Analysis, LRG; Investigation, LRG, LU, FA and PNM; Writing – Original Draft, LRG and PNM; Writing – Review & Editing, LRG, RI, LU, FA, HK, CET, BC, LJP and PNM; Visualization, LRG; Funding Acquisition, LJP and PNM; Supervision, LJP and PNM.

## Acknowledgments

This study was funded by grants from Medical Research Council (MR/S004688/1; https://mrc.ukri.org/) and The Humane Research Trust (145355; https://www.humaneresearch.org.uk/). The funders had no role in study design, data collection and analysis, decision to publish, or preparation of the manuscript. For the purpose of open access, the author has applied a Creative Commons Attribution (CC BY) licence to any Author Accepted Manuscript version arising.

The authors would like to acknowledge the Wolfson Light Microscopy Facility at The University of Sheffield and in particular Dr. Christa Walther for her help and advice with the fluorescent peptide analysis. Furthermore, the authors would like to acknowledge Prof. Craig Murdoch and Dr. Helen Colley for providing access to the primary normal human tonsillar keratinocytes.

