## Supplemental Data for "CD9 co-operation with syndecan-1 is required for a major staphylococcal adhesion pathway"

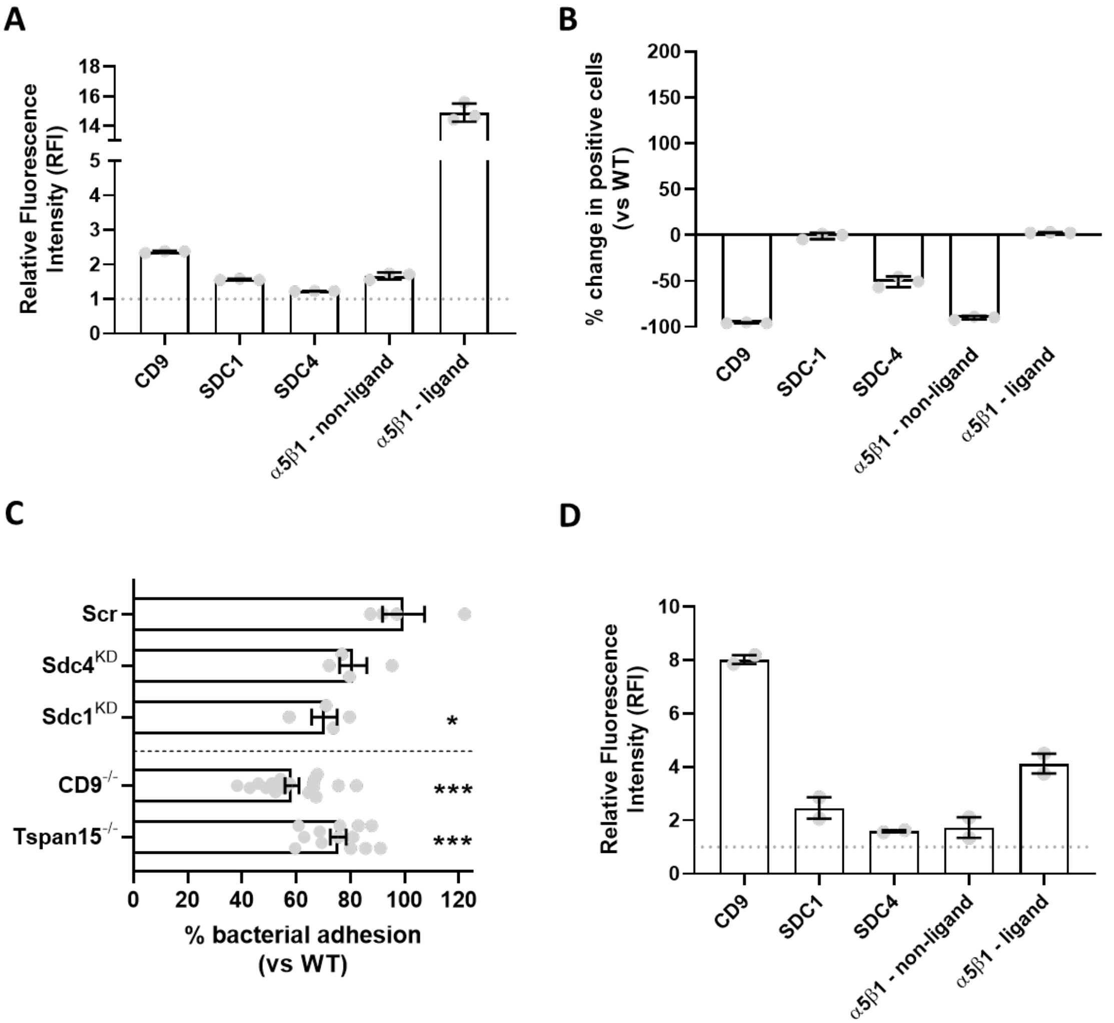


**Fig. S1. Knockout or knockdown of CD9 and putative staphylococcal receptors reduces *S. aureus* adhesion to A549 cells.** (A) WT cells were treated with an anti-CD9 antibody (602.29), an anti-SDC-1 antibody (B-A38), an anti-SDC-4 antibody (5G9), an anti-α5 antibody (JBS5) or an anti-β1 antibody (AIIB2). Cell surface expression was determined using a FITC-conjugated secondary antibody. Relative fluorescence was determined by dividing the test antibody by the isotype control. (B) Cell surface expression of staphylococcal receptors differ in knockout cells. WT or $CD9^{-/-}$ cells were treated with an anti-CD9 antibody (602.29), an anti-SDC-1 antibody (B-A38), an anti-SDC-4 antibody (5G9), an anti-α5 antibody (JBS5) or an anti-β1 antibody (AIIB2). Cell surface expression was determined using a FITC-conjugated secondary antibody. $CD9^{-/-}$ cells were compared against the wild-type cells to give the % change. (C) Knockout or shRNA knockdown cells were infected with SH1000 for 60 mins at an MOI=50. (D) NTKs were treated with an anti-CD9 antibody (602.29), an anti-SDC-1 antibody (B-A38), an anti-SDC-4 antibody (5G9), an anti-α5 antibody (JBS5) or an anti-β1 antibody (AIIB2). Cell surface expression was determined using a FITC-conjugated secondary antibody. Relative fluorescence was determined by dividing the test antibody by the isotype control. n$\geq$2, mean $\pm$ SEM, One-Way ANOVA.

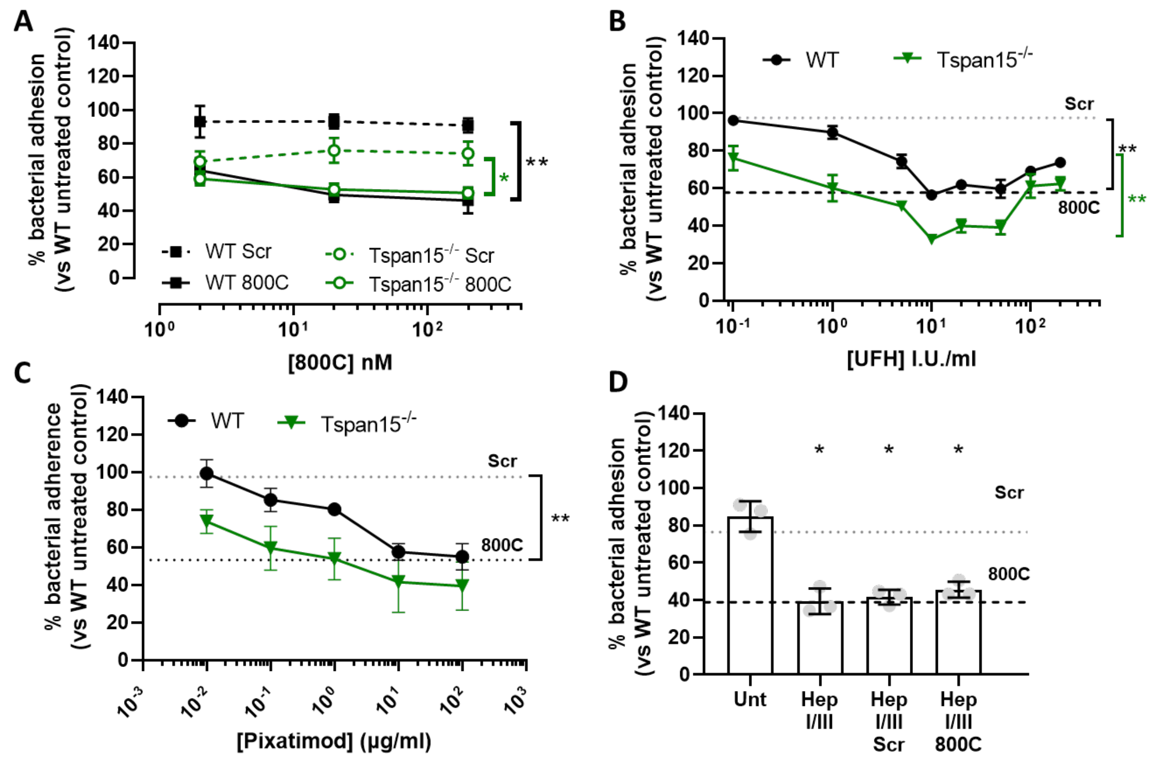


**Fig. S2. CD9-derived peptides, heparin analogues and removal of heparan sulphates reduce staphylococcal adherence in Tspan15$^{-/-}$ cells.** Tspan15$^{-/-}$ cells were infected with SH1000 for 60 mins at an MOI=50. (A) WT (black) or Tspan15$^{-/-}$ (green) cells treated with scrambled (dotted) or 800C peptide (solid) for 60 mins prior to infection. WT (black) or Tspan15$^{-/-}$ (green) cells treated with UFH (B), or various concentrations of pixatimod (C). Tspan15$^{-/-}$ cells were treated with 0.5U/ml heparinase I/III for 3 hours prior to infection (D). Effect of CD9-derived peptide treatment (200nM) shown by dotted lines (B-D). n$\geq$3, mean $\pm$ SEM, One-Way ANOVA.

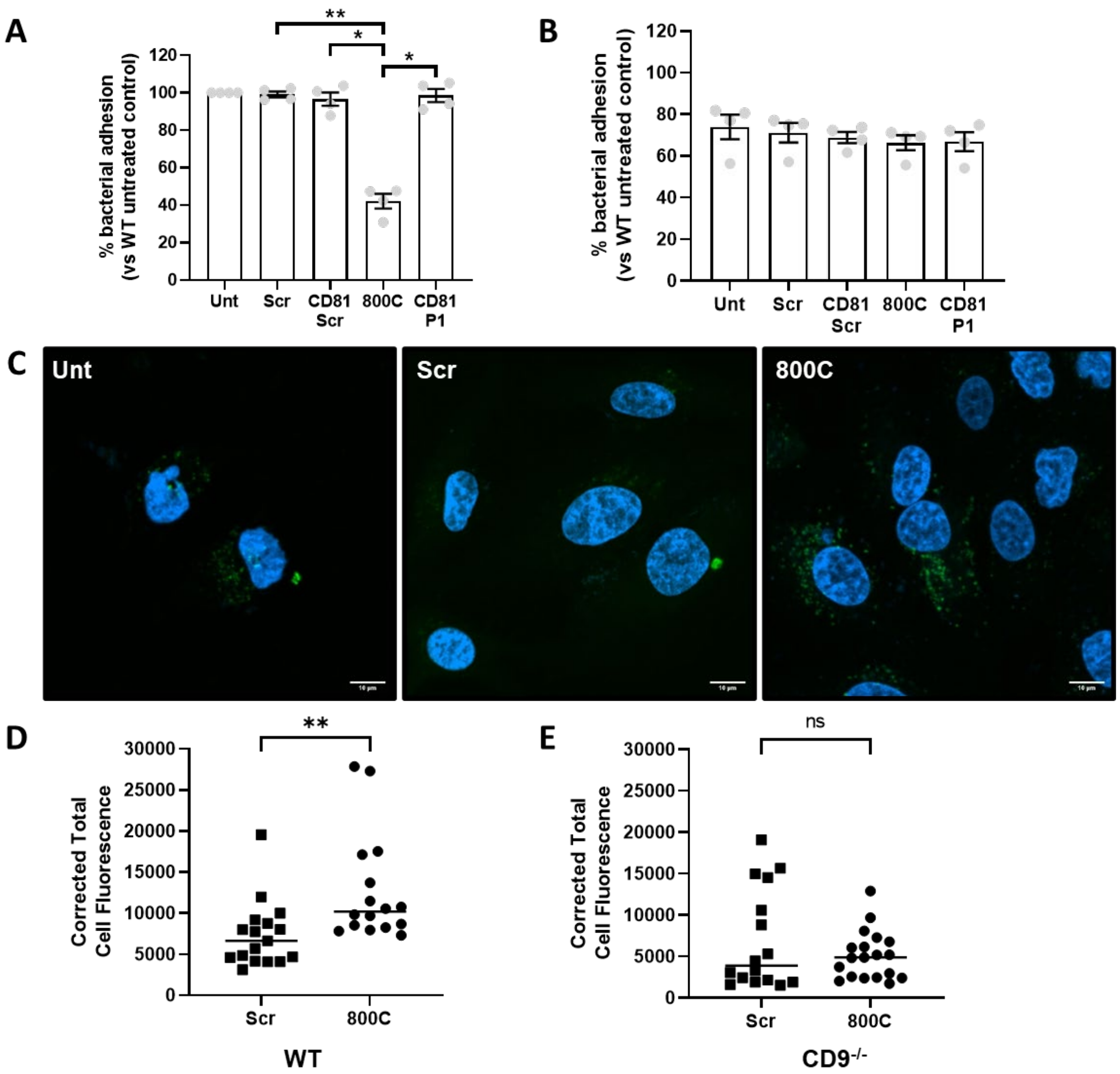


**Fig. S3. 800C interacts with cells expressing CD9 and specifically inhibits staphylococcal adherence.** WT (A) or CD9$^{-/-}$ (B) cells were infected with SH1000 for 60 mins at an MOI=50. Cells were treated with 800C, CD81 P1 or their scrambled counterparts for 60 mins prior to infection. n=4, mean ± SEM, One-Way ANOVA. WT cells were treated with FAM-tagged 800C or scrambled peptide for 30 minutes. Cells were fixed with paraformaldehyde and imaged by confocal microscopy. Representative images are shown in panel C. Corrected total cell fluorescence was calculated for individual WT cells (D) or CD9$^{-/-}$ cells (E) treated with either FAM-tagged scrambled or 800C peptide. n≥16, median, data was analysed by Mann-Whitney test.

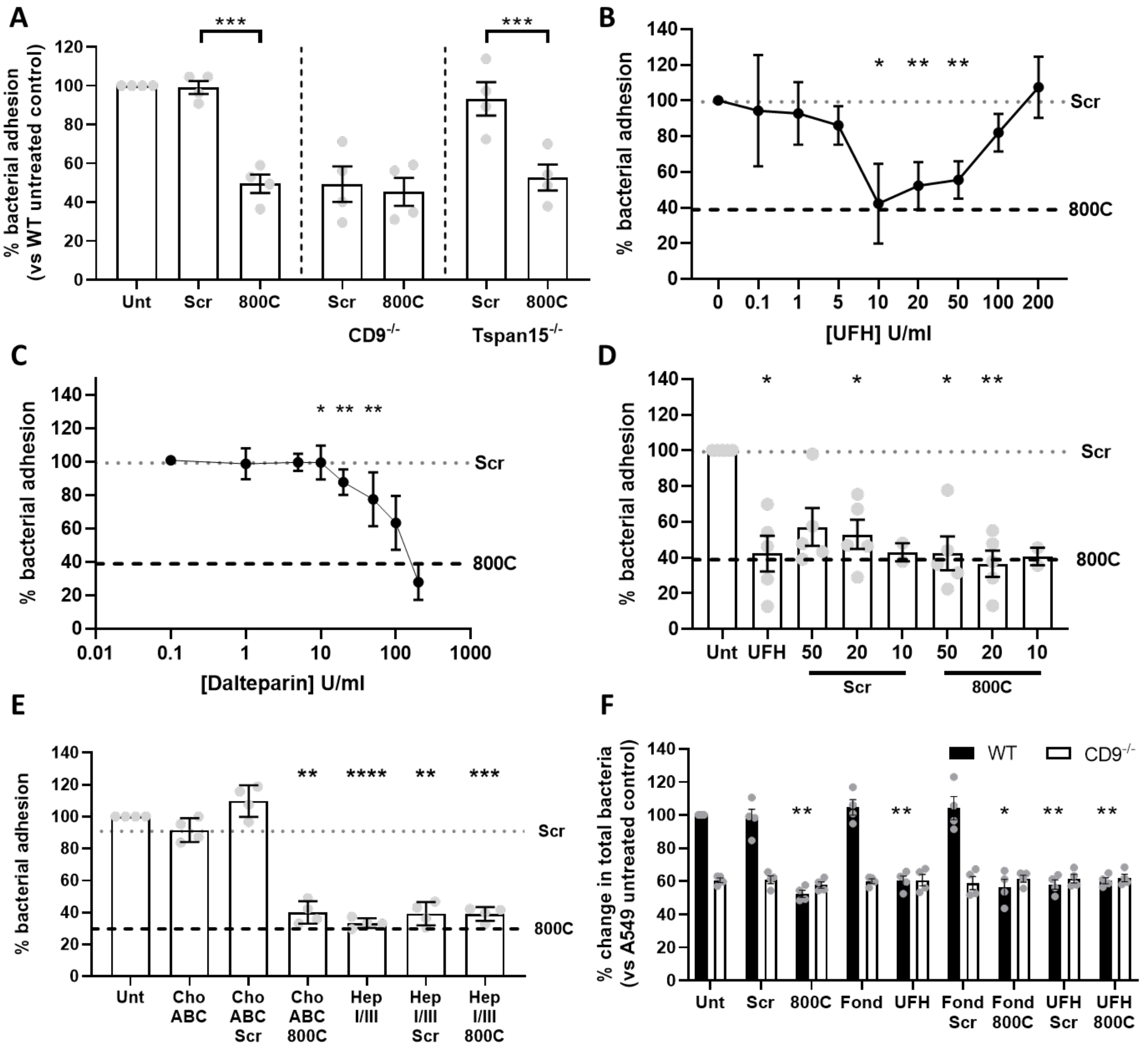


**Fig. S4. CD9-derived peptides can affect clinical *S. aureus* isolates and reduce staphylococcal adhesion to human keratinocytes similar to heparin analogue treatments.** (A) WT, CD9$^{-/-}$ or TSPAN15$^{-/-}$ cells were infected with MRSA1 for 60 mins at an MOI=50. Cells were treated with scrambled or 800C peptide for 60 mins prior to infection. (B-E) HaCaT cells were infected with SH1000 for 60 mins at an MOI=50. (B) HaCaT cells treated with UFH for 60 mins prior to infection. (C) HaCaT cells treated with dalteparin. (D) Cells were treated with a combination of various concentrations of UFH with either scrambled or 800C peptide. (E) Cells were treated with 0.25U/ml chondroitinase ABC or 0.5U/ml heparinase I/III for 3 hours prior to infection. Effect of CD9-derived peptide treatment (200nM) shown by dotted lines (A-E). (F) Enumeration of the total number of bacteria adhered to epithelial cells by microscopy. WT or CD9$^{-/-}$ cells on glass coverslips were treated with 800C (200nM), UFH (10U/ml), Fondaparinux (10µg/ml) or various combinations of these treatments and infected with SH1000 for 60 minutes at an MOI=50. Coverslips were stained with Giemsa stain and examined by light microscopy. The total number of bacteria associated to 100 cells was enumerated and the fold change calculated by comparing against the WT untreated control. n≥3, mean ± SEM, One-Way ANOVA.

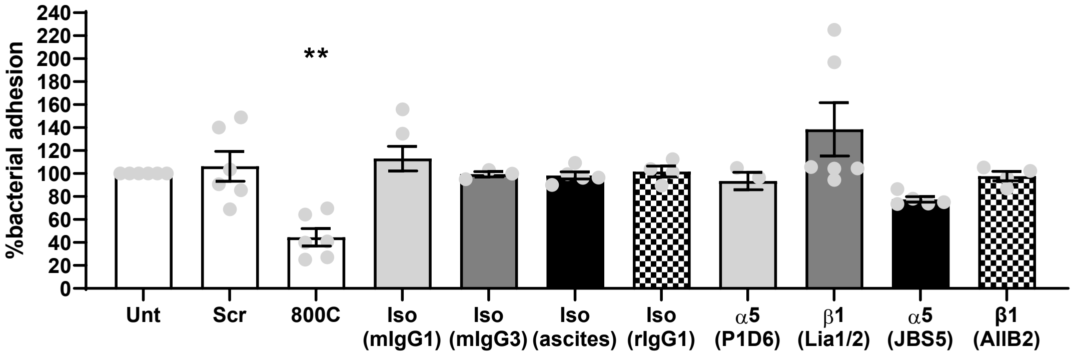


**Fig. S5. Various anti-α5β1 antibodies have little effect on staphylococcal adherence to human keratinocytes.** HaCaT cells were infected with SH1000 for 60 mins at an MOI=50. Cells were treated with scrambled or 800C peptide, anti-α5 antibodies (P1D6 and JBS5), anti-β1 antibodies (Lia1/2 or AIIB2), or appropriate isotype controls for 60 mins prior to infection. n≥4, mean ± SEM One-Way ANOVA.

**Table S1. Reduction of staphylococcal adhesion using either CD9-derived peptides or unfractionated heparin in various cell lines.**

| Cell line | Untreated cfu | 800C (200nM) cfu (% reduction) | UFH (10 U/ml) cfu (% reduction) |
|---|---|---|---|
| A549 | 27285.7 ± 6651 | 13952.4 ± 5932.9 (49.3 ± 13.7) | 12857.1 ± 4802.8 (52.3 ± 13.4) |
| HaCaT | 25555.6 ± 5787.2 | 8500 ± 3044.3 (64.8 ± 16.2) | 9740.7 ± 3966.3 (59.5 ± 20.4) |
| HCE-2 | 16250 ± 5755 | 7958.3 ± 2916.7 (50 ± 8.65) | 5375 ± 1945.4 (63.9 ± 15.8) |
| IPEC-J2 | 19555.6 ± 1873.3 | 8666.7 ± 4419 (56.8 ± 18.6) | 6055.6 ± 1134.4 (69 ± 5.1) |
| NTK | 47333.3 ± 5897.3 | 23388.9 ± 7743 (51.4 ± 9.9) | 24888.9 ± 9229.1 (48.4 ± 12.5) |

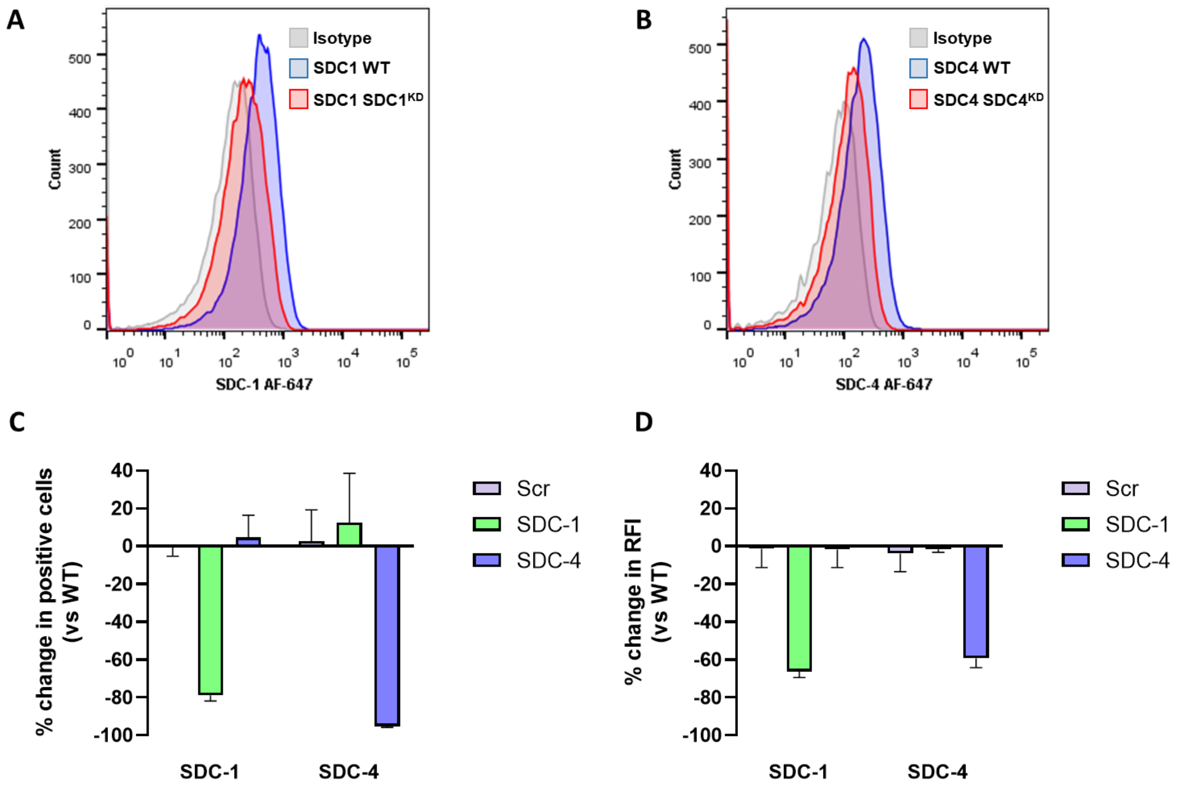


**Fig. S6. Confirmation of SDC knockdown in A549 cells with shRNA.** shRNA lentiviral infected WT cells were detached and fixed in methanol. Fixed cells were treated with an AlexaFluor-647 anti-SDC-1 antibody (B-A38; A) or an anti-SDC-4 antibody (5G9; B) to determine cell expression. Representative histograms are shown in A and B. Relative fluorescence was determined by subtracting the isotype control from the test antibody. Knockdown cells were compared against the WT cells to give the % change. n=3, mean ± SEM.

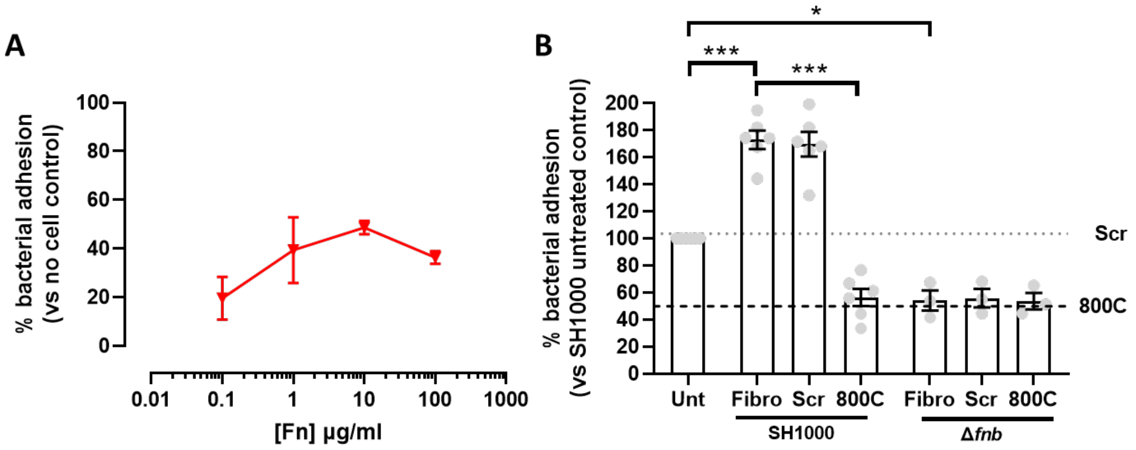


**Fig. S7. Non-specific bacterial binding increases after treatment with exogenous Fn but does not increase binding of a Δ*fnb* mutant.** (A) 96 well plates were treated with various concentrations of Fn for one hour prior to infection. Treated wells were challenged with SH1000 for 60 minutes with bacterial numbers comparable to cellular infection. The number of colonies was compared to those of untreated wells to give the % change. (B) Fn was added to WT cells in combination with 800C or a scrambled peptide for one hour prior to infection with either SH1000 or SH1000 Δ*fnb*. n=3, mean ± SEM.
